# Bead-free deterministic DNA barcoding using vacuum-driven loading of aqueous oligonucleotides to microwell arrays

**DOI:** 10.1101/2025.10.29.684720

**Authors:** Patrycja Baranowska, Trinh Lam, Amy E. Herr

## Abstract

Achieving high throughput in experiments requiring sample indexing depends primarily on precise and reproducible reagent deposition prior to analysis. Contemporary droplet and microwell systems utilize random deposition of oligonucleotide-coated beads into reaction chambers, requiring costly bead synthesis and offering limited control over the final distribution of barcoded beads. As an alternative, we present deterministic, aqueous barcoding of 512 arrayed microwells using a multi-layer, vacuum-driven microfluidic network. To uniquely barcode each of the 512 microwells, we deposit DNA oligonucleotide solutions designed using a Combinatorial Dual Indexing (CDI) (i5, i7) scheme via deterministic loading. Deterministic fluid loading is achieved by sequentially mating two bifurcated, orthogonal microchannel networks to a planar microwell array. The microchannel networks actuate fluid flow through a combination of an applied vacuum force and a dead-end channel design. After loading the oligonucleotide solutions, we observed uniform barcode patterning across the arrays of microwells (∼20% CV), reasonable barcode loading times (30 - 40 min per step), and reduced reagent use (∼ 8-16 µL at 25 µM oligos vs. 10-50 µL at 100 µM for bead systems). We detected cross-contamination in ∼4% of the microwells. Following DNA barcode delivery, on-chip PCR of nuclear DNA from a breast-cancer cell line having the characteristics of the differentiated mammary epithelium (MCF7) was successfully performed, and off-chip quality control of the amplified breast cancer DNA was completed. Overall, we describe a deterministic and bead-free DNA barcoding strategy for efficient barcoding of microwell arrays that are important in single-cell analyses.

## Introduction

The rapid development of high-throughput systems in fields such as molecular diagnostics, transcriptomics and genomics has created a growing demand for precise and scalable methods for spatially controlled reagent delivery.^1–3^ The accuracy and reproducibility of microscale fluid delivery are key factors in data quality. Consequently, there is a strong need for deterministic, spatially patterned reagent deposition systems offering a minimal risk of cross-contamination and maximum operational stability. Microfluidic systems provide an efficient and precise platform for the controlled manipulation of fluids at the microscale, and are thus well suited for delivery of metered reagent volumes in structured receiving geometries (e.g., microwells).^4^ Applications that require high throughput and precise sample tracking benefit, such as barcoding experiments preparing samples for analysis on modern next-generation sequencing (NGS) platforms.^5^

Bead-based barcoding is a cornerstone barcoding techique^6–9^ that is frequently implemented in one of two compartment geometries: droplets and microwells. In droplet-based approaches, individual cells or nuclei are encapsulated within microdroplets containing barcoded beads and lysis buffer (e.g., 10x Genomics Chromium^10^, Drop-seq^1^, inDrops^11^). In microwell or plate-based approaches (e.g., Smart-seq2^12^, Seq-Well^13^, sci-RNA-seq^14^), cells are co-deposited into wells with magnetic beads that are coated with unique barcodes. Upon cell lysis in the microwell, RNA or DNA binds to the bead, thus precisely assigning cellular transcripts of interest to the individual originating cell. The magnetic properties of the beads allow for convenient isolation and downstream processing.^15^ Despite advantages and widespread use, bead-based methods have limitations (i.e., random assignment of barcodes, high costs, bead aggregation, additional steps in barcode preparation, and loss of material during magnetic separation) some of which may be exacerbated by the application of a magnetic field.^16–19^

In contrast to bead-based (heterogeneous) barcoding approaches, spatial patterning of aqueous barcodes (homogeneous) relies primarily on the precision of a dispensing fluidic system (e.g., microchannel architecture, sealing, and flow control). Aqueous barcodes offer greater flexibility in barcode design and facilitate combinatorial indexing to generate a unique combination of nucleic acid barcodes for each cell, although aqueous barcodes require precise control of fluid delivery to prevent barcode cross-contamination (i.e., diffusion, mixing).^20^ The performance of such systems therefore depends on the mechanical design, channel alignment, seal integrity, and resistance to diffusion or unintended mixing. One of the most well-known and proven implementations of spatial reagent delivery has been demonstrated in deterministic barcoding methods such as DBiT-seq^21,22^, where parallel microfluidic channels deliver two sets of barcodes to tissue sections, resulting in grid-patterning of uniquely coded pixels across each tissue slice for spatial transcriptomics. Furthermore, this technique offers many possibilities for extension and modification, as demonstrated by its refinement - xDBiT^23^, which supports the simultaneous analysis of multiple tissue sections while maintaining high spatial resolution. Rather than focusing on sequencing chemistry, such systems highlight the importance of channel uniformity, leak-free interfacing, reproducible alignment, and reliable fluid exchange across repeated runs.

Inspired by fluidic grid patterning^24,25^ and CDI deterministic barcoding, we sequentially apply two vacuum-driven orthogonally oriented microchannel layers (horizontal and vertical) to pattern a unique aqueous DNA barcode in each microwell of a 512 microwell array. Unlike previously described systems^26–28^, vacuum is generated by facile interfacing each dispensing microfluidic chip to suction – at a single-point – from a house vacuum system. Importantly, we design the deterministic approach using aqueous DNA barcodes to eliminate resource-intensive synthesis and functionalization of beads with oligonucleotides. Concomitantly, aqueous barcode formulations have the potential to reduce reagent consumption compared to existing protocols, thus reducing operational costs and biological material loss.^29^ Here, we sought to harness the precision flow control afforded by microsystems to reduce microwell-to-microwell barcode cross-contamination. By combining high spatial resolution with operational simplicity and low instrumentation requirements, our platform offers an attractive and affordable alternative to current DNA barcoding techniques.^30^

## Experimental Section

### Choice of aqueous barcodes for combinatorial dual indexing of the microwell array

A combinatorial dual indexing (CDI) scheme was employed in conjunction with a deterministic barcode loading strategy to pattern each microwell of the 512 microwell array with a unique barcode index. We used Illumina i5 and i7 indexes as solution-phase barcodes. A 25-µM solution of the i5 barcode (Integrated DNA Technologies; in ultrapure water) was used for the horizontal loading step. Similarly, a 25-µM solution of the i7 barcode (Integrated DNA Technologies; in ultrapure water) was used as the barcode for the vertical loading step. The barcode sequences are as follows:

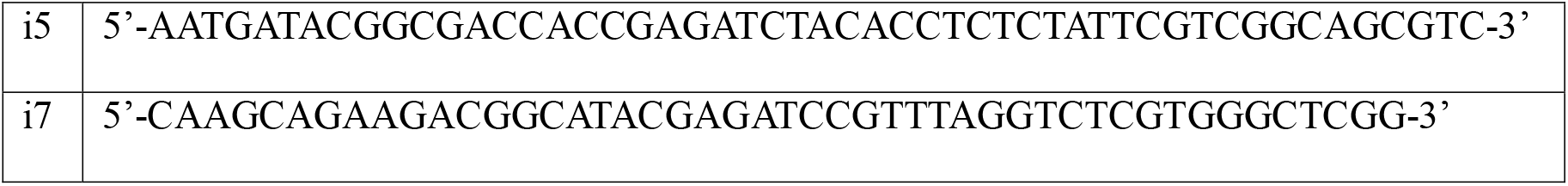

### Device geometry and fabrication

The deterministic barcode loading strategy uses five PDMS-device layers: a 512 microwell array, a vacuum network, and three layers each consisting of different bifurcated microfluidic channel networks: a horizontal barcode delivery network, a vertical barcode delivery network, and a PCR reagent delivery network. Soft lithography was used to fabricate all layers: SU-8 3500 (Kayaku Advanced Materials) for horizontal, vertical, and PCR delivery layers, and SU-8 2100 for microwell and vacuum layers. Spin coating, soft baking, and PDMS development followed standard protocols (see SI). After each PDMS layer was cast, inlet holes were punched in the delivery layers (∅ = 1.5 mm) and the vacuum layer (∅ = 1 mm) using a biopsy punch. The vacuum layer had 4 ports connected to the house vacuum; the horizontal, vertical, and PCR layers had 16, 32, and 1 open ports, respectively.

### Assembly of the multi-layer device and fluid routing for deterministic barcode loading

To assess microwell filling, we employed both brightfield and fluorescence imaging to visualize the microchannel-flow and microwell-filling characteristics. Imaging-based cross-contamination assessment used progressively more stringent test samples, progressing from solutions of colorimetric (food) dyes, to fluorescein-o-acrylate (FITC acrylate, 1mg/ml in DMSO, Sigma-Aldrich), to finally solutions of oligonucleotides conjugated with fluorophores (biotin-i51-488 and oligo-A549, 25 uM in ultrapure water, Integrated DNA Technologies, IDT).

The microfluidic device comprising a cleaned glass slide, a PDMS microwell layer, and experiment-specific delivery and vacuum layers was plasma-activated, aligned, and temporarily sealed under vacuum before loading the barcode solution into the inlets. After fluid loading, the vacuum layer was removed, the chip was secured with PCR film, and centrifuged at 3000 RPM for 2 min, with full procedural details provided in the Supplementary Information.

### Cross-contamination during deterministic loading of the microwell array

To assess potential cross-contamination between adjacent microwells, two solutions of oligonucleotide, with each oligonucleotide conjugated with distinct fluorescent tag, were prepared: oligo-A549 (Integrated DNA Technologies, IDT) and biotin-i51-488 (Integrated DNA Technologies, IDT) both in concentrations 25 uM prepared in ultrapure water. Biotin-i51-488 was used for the first barcoding step (horizontal) and oligo-A549 was used for the second barcoding step (vertical). The first barcoding step was performed by flowing Biotin-i51-488 solution to every second microchannel of the microsystem (channel pattern: oligo, water, oligo, water, etc.). In this way, horizontal rows of microwells were patterned with alternating solutions of fluorescent oligos and clear water. Fluid was introduced using a 20 μl pipette, followed by a 30-40 min loading duration. This alternating pattern allows ready visualization of any leakage of fluorescent solution between adjacent microwell rows. After all of the microwells were filled with the respective fluid, the vacuum layer was carefully removed, the microwell and inlet areas were sealed with q225 PCR sealing film (Quantagene, KUBO) and the system was centrifuged at 3000 RPM for 2 min. Fluorescence intensity in both the 488-nm and 549-nm spectral channels was measured across the microwell array.

Then, to prevent cross-contamination between microwells, the solution of oligonucleotides in the microwells were dried in an oven at 35°C for 15 min. The fluidic layer was then carefully removed, and the microwells were further dried at 35°C for 20 min. Next, the vertical delivery layer was aligned to the microwell array layer containing the dried oligonucleotides. The vacuum layer was applied on top of the assembly, connected to the vacuum tubing, and all inlets were sealed with tape (600 Scotch Crystal, 3M). Vacuum was applied for ∼1 min. Then the inlets were opened and oligo-A549 barcode solution was pipetted into to every second inlet (channel pattern: oligo, water, oligo, water, etc.) using a 20 μl pipette to achieve the desired alternating barcode pattern in the microwell rows. The system equilibrated for 30–40 min, with visual monitoring of the loading process. After loading, the vacuum layer was removed, and the system was resealed with q225 PCR film (non-sticky side, Quantagene, KUBO). The chip was centrifuged again at 3000 RPM for 2 min. Fluorescence intensity in both the 488-nm and 549-nm spectral channels was again measured across the microwell array.

### Isolation of nuclei from the MCF-7 mammalian cell line

Nuclei were isolated using ATAC-RSB buffer which was prepared by mixing 500 µL of 1M Tris-HCl pH 7.4 (10 mM), 100 µL of 5M NaCl (10 mM), 150 µL of 1M MgCl2 (3 mM), and 49.25 mL of molecular biology grade water. The lysis buffer was prepared by adding 0.1% IGEPAL CA-630, 0.1% Tween-20, and 0.01% digitonin to ATAC-RSB buffer to reach the final volume. The wash buffer contained 0.1% Tween-20 in ATAC-RSB buffer.

One million viable MCF7 cells were transferred to 1.5 mL LoBind Eppendorf tubes and pelleted at 500 × g for 5 min at 4 °C. After discarding the supernatant, the pellet was rinsed in 1 mL ice-cold PBS, spun again under the same conditions, and the PBS removed. Cells were then lysed by adding 300 µL chilled lysis buffer, mixing gently, and incubating on ice for 5 min. Lysis was quenched with 1 mL cold wash buffer, followed by five gentle inversions. Nuclei were collected with a two-step spin - first with the hinges facing in, then facing out - both at 500 × g for 3 min at 4 °C. The supernatant was aspirated, and nuclei were resuspended in 250 µL of wash buffer. Nuclei were resuspended in 250 µL wash buffer using a wide-bore tip (Rainin) and counted on a Countess after mixing 1:1 with Trypan Blue. Then 50,000 nuclei were added to 1 ml of cold wash buffer and centrifuged first with the hinges facing in, then out - both at 500 × g for 5 min at 4 °C. The supernatant was removed and nuclei pellet was resuspended in 50 µl tagmentation cocktail prepared using 1× tagmentation buffer (Illumina Inc), 0.1% Tween 20, and 0.01% digitonin (Promega). TDE1 tagmentation enzyme (Tn5, Illumina Inc) was added to the cocktail at a ratio of 1:15. The tagmentation reaction was incubated at 37°C on a thermoshaker at 1000 RPM for 30 min. After tagmentation, 250 µL DNA binding buffer from Zymo DNA Clean and Concentrator Kit was added to the reaction, followed by clean up using the same kit. The concentration of tagmented DNA was measured by fluorimetry (Qubit).

### PCR-on-chip

To prepare the chip for on-chip PCR and amplification, alternating rows of microwells were patterned with an solution-phase i5 index (via horizontal delivery layer) or an solution-phase i7 index (via vertical delivery layer). After barcoding, the microwell array layer was aligned with the PCR delivery layer and the vacuum layer was mated on top. The vacuum layer had four ports connected to the internal vacuum and one inlet for introducing PCR reagents. Next, the PCR master mix, composed of ultrapure water (Invitrogen), 1× NEB Master Mix (New England Biolabs), 0.005% Tween-20 (Promega), and BSA (1□mg/ml; Sigma-Aldrich) was prepared, and 8□μl of the master mix was combined with 2□μl of tagmented MCF7 DNA. Next, as previously described, the inlet was sealed with tape (600 Scotch Crystal, 3M), vacuum was applied for less <1 min, the inlet was opened, and the PCR mastermix with tagmented DNA (concentration: 0.454ng/ml) was introduced into the system. After confirming proper loading and stable applied vacuum, 20 μL of mineral oil (Sigma Aldrich) was added, followed by an additional 50 μL of mineral oil on top of the inlet. Vacuum was released (the tubes were removed) and water was loaded into the vacuum layer using a syringe and all four ports to the vacuum. After that, all the ports were sealed with thermally conductive substrate polyimide film (Kapton Tape, thickness = 0.069 mm, 3M). Next, the microfluidic chip was placed in the Proflex PCR system (Life Technologies), covered with a coverslip, sealed, and subjected to 15 amplification cycles (98°C for 10 s, 63°C for 30 s, 72°C for 60 s) after an initial denaturation at 98°C for 30 s and gap filling at 72°C for 5 min.

### DNA collection and cleanup

Post-PCR, the vacuum layer was carefully removed, and the layer and microwell array were cut and transferred to a petri dish. A series of washes to the delivery layer and microwell were performed using 2600 μL DNA Binding Buffer (Zymo Research), and the reagents were collected via a laboratory vacuum system. DNA was cleaned using Zymo DNA Clean and Concentrator Kit as instructed. Final elution was performed with 7 μL Elution Buffer (Zymo Research).

### DNA quantification and quality control

DNA concentration was measured using the Qubit fluorometry system according to the manufacturer’s instructions for the Qubit™ dsDNA HS Reagent (Invitrogen). DNA electropherograms were generated and assessed using a TapeStation (Agilent Technologies) with High Sensitivity D1000 Reagents.

### Brightfield and fluorescence microscopy

Fluorescence intensity was measured using an Olympus IX51 microscope and filter sets for GFP and Texas Red equipped with ImageJ (NIH, USA) imaging software.

## Results

### Barcoding principle and design of deterministic loading process

To design a bead-free barcoding approach suitable for a 512 microwell array (Figure 1A, B), we designed a barcoding strategy with two attributes: (1) a CDI scheme is employed for design of the DNA oligonucleotide barcodes using Illumina i5 and i7 indexes and (2) a deterministic barcode-loading scheme fluidically delivers the solution-phase DNA oligonucleotide barcode combinations to each of the 512 microwells in the planar microwell array. The barcode scheme is designed for endpoint Illumina sequencing of pooled barcoded DNA samples, with an interest here in the intermediate steps of on-chip PCR followed by DNA quality control by Tapestation analysis (Figure 1C). Adoption of a bead-free, aqueous barcoding strategy requires low-crosstalk delivery of unique barcodes to each of the 512 microwells. Informed by the work of Hu et al.^24^ we designed a vacuum-driven microfluidic system based on a grid-patterning approach inspired by DBiT-Seq^21^. To uniquely barcode each microwell, we used a multi-step, vacuum-assisted loading process. Unlike other approaches^31,32^, fluid flow was driven by house vacuum via pressure differences across the microwell, delivery, and vacuum layers.^33^ The first barcode set was delivered via the horizontal delivery network (i5) and then a second set via the vertical delivery network (i7), creating a CDI scheme.

**Figure 1.**
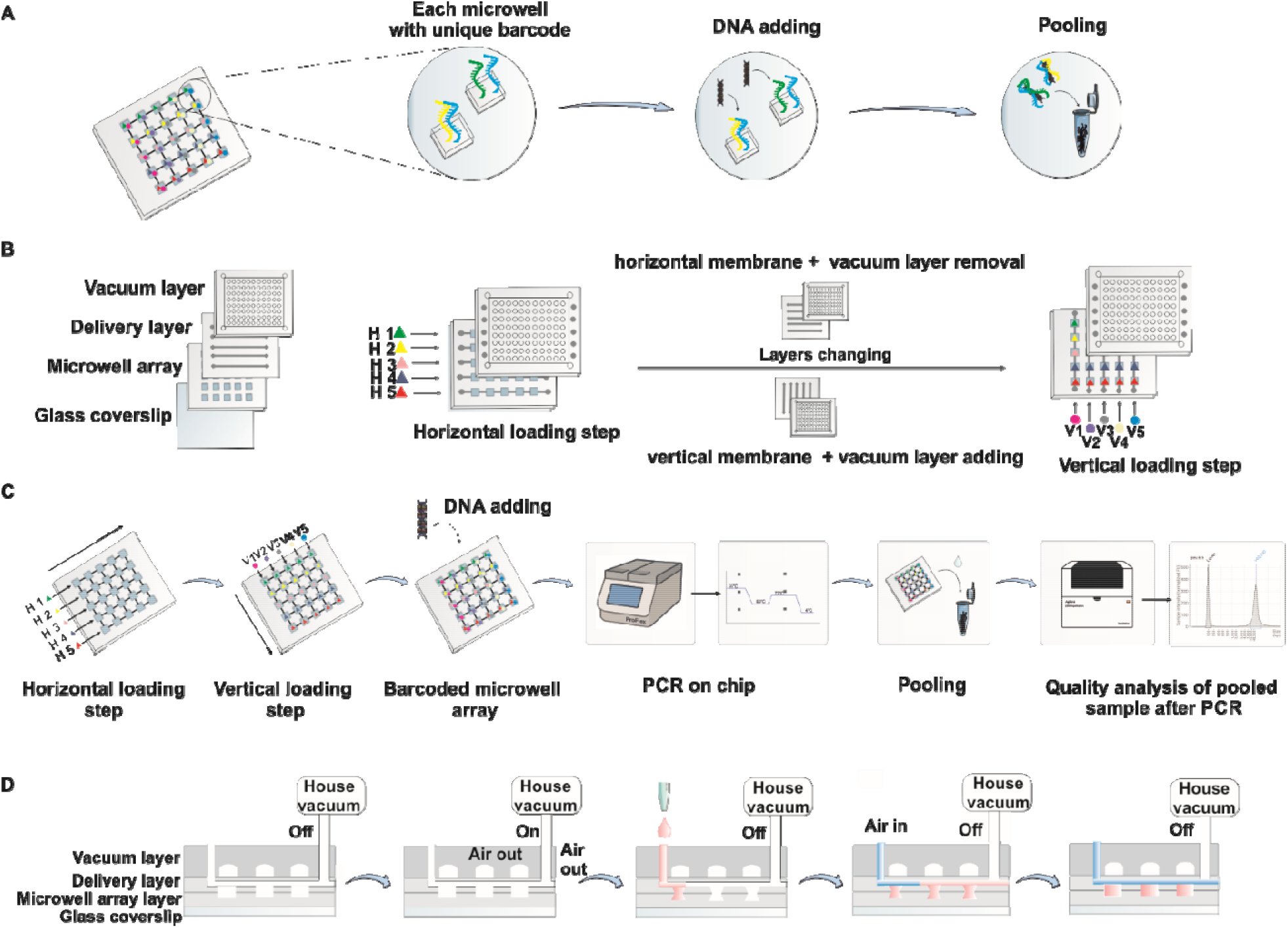
Deterministic, bead-free oligonucleotide barcoding of arrayed microwells using a multi-layer vacuum-driven microfluidic network. (A) Use of aqueous barcodes and deterministic loading of unique barcodes to each microwell in the array allows pooling of barcoded samples for downstream analysis (e.g., sequencing). (B) Diagram of the barcoding system. The process includes sequential horizontal and vertical loading stages, with drying and layer exchange between the steps to ensure deterministic assignment of barcodes. (C) To achieve 512 uniquely barcoded microwells, the workflow includes a two-step aqueous barcode loading process (horizontal and the vertical delivery of barcodes), sample (e.g., DNA) loading, on-chip PCR, collection and pooling of barcoded samples for pooled analysis of DNA quality by electrophoresis (Tapestation). (D) A multi-layer PDMS microdevice is designed to pattern unique aqueous barcodes into each of an array of dead-end microwells using the gas permeability of PDMS and applied suction to drive the flow of reagents to each microwell.

This approach produces 512 microwells, each with a unique barcode combination. The microwell, delivery, and vacuum layers were manually aligned under a stereomicroscope, sealed, and connected to house vacuum (45-60 s). Evacuation through the PDMS layer created a pressure gradient (Figure 1D), confirmed by microwell perimeter bowing. After releasing the vacuum, barcode solutions (0.3-0.7 µm per channel) were pipetted into inlets, flowing by suction through bifurcated microchannels to fill dead-end microwells. Air was introduced to clear excess solution and cap each microwell, enabling deterministic barcoding and subsequent on-chip PCR. The process followed two steps: first, the vertical network delivered i5 solutions, which evaporated to leave residues in the patterned microwells; next, the vertical layer was removed and replaced with a perpendicular horizontal network for i7 delivery. After a second loading of the DNA oligonucleotide solution, each microwell was anticipated to contain a unique i5/i7 combination.

### Microfluidic device designs for deterministic barcoding process

The vacuum-assisted microfluidic system consists of three PDMS layers: a microwell array layer, a set of three interchangeable fluid delivery layers (Figure 2 A) (used sequentially for horizontal barcode loading, vertical barcode loading, and PCR reagent delivery), and a vacuum layer. The microwell array layer contains only microwells (512 in total, 32 × 16) without channels, while the fluidic networks, inlets, and narrow necks (90□µm × 150□µm) are fully integrated into the 100□µm-thick delivery layers. The narrow neck functions as a microfluidic constriction that reduces or prevents backflow from the microwell into the main channel^34,35^, thereby reducing the risk of cross-contamination between adjacent microwells (Figure 2 B).

**Figure 2.**
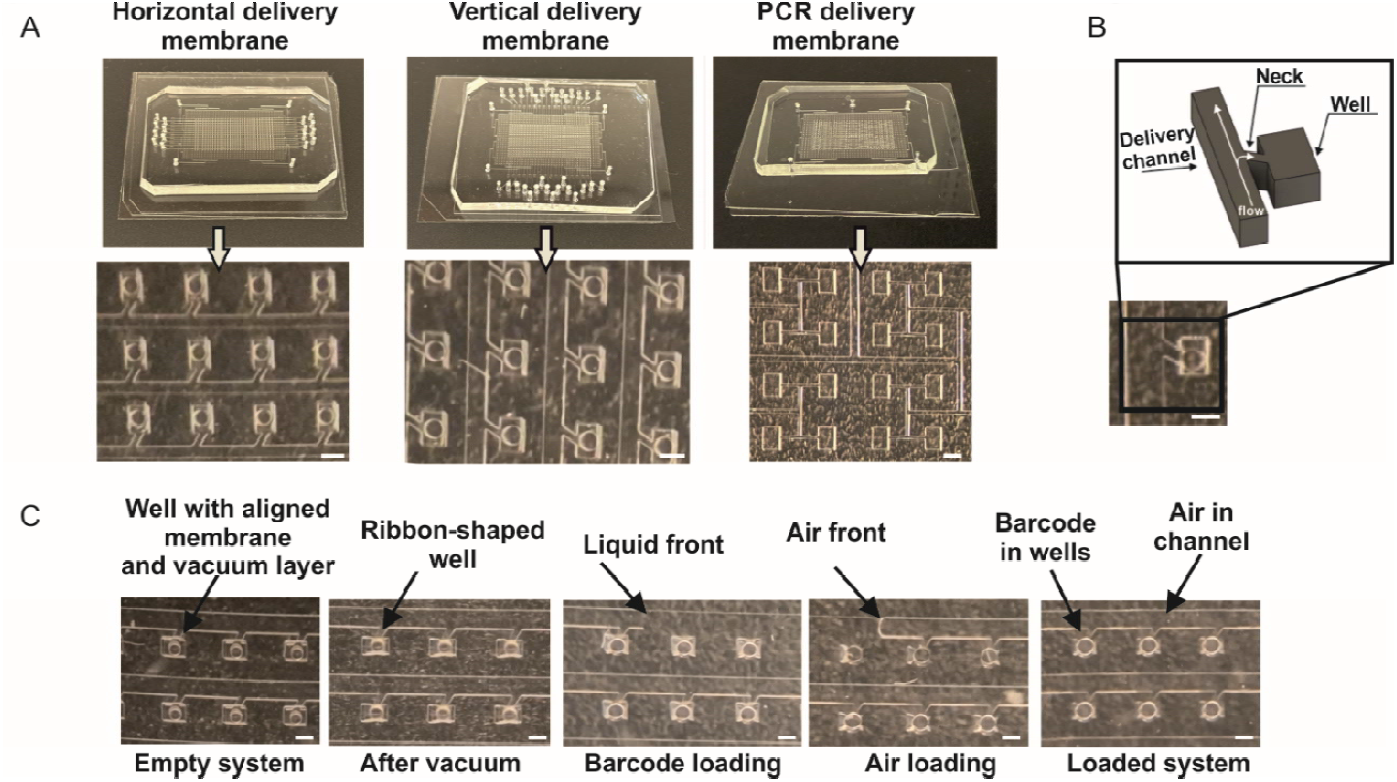
Microfluidic networks are designed to pattern specific microwells with aqueous oligonucleotide barcode solutions. (A) Brightfield images and micrographs show three different microfluidic network device configurations, with discrete functions including: (1) a horizontal delivery layer, (2) a vertical delivery layer, and (3) a PCR-reagent delivery layer. Each microfluidic network layer is mated to the microwell array layer which is seated on a rigid glass slide. (B) Bright-field micrograph and schematic representative of the interface between the main delivery microchannel and a necked microwell. (C) Brightfield micrographs show the sequential stages of barcode-solution loading using the delivery layers to fill each microwell. Scale bars: 250 µm.

Each microwell in the microwell array layer measures 250 × 350 × 100□µm, with horizontal and vertical pitches of 550□µm and 450□µm to reduce the risk of cross-contamination. When mated to the fluid delivery layers, the assembly extends the effective microwell depth to 200□µm. Each microchannel has a separate inlet to confine the introduction of barcode solutions uniquely to that microchannel and the associated row of microwells.^24^ Unlike bead-based barcoding systems, the aqueous barcoding strategy using fluid filling and not external magnetic forces (e.g., as used with the more common oligo-conjugated magnetic beads) to retain barcodes inside microwells after loading (Figure 2 C), making the constriction design critical for robust operation.^13,36^

During the horizontal loading step, i5 barcodes are delivered through 16 parallel channels (45□mm × 150□µm × 100□µm) with 1.5□mm inlets. The vertical delivery layer introduces i7 barcodes via 32 perpendicular channels (34□mm × 200□µm × 100□µm), each with a dead-end design, allowing deterministic CDI while minimizing uncontrolled flow. The vacuum layer (∼7□mm thick) contains four ports for facile connection to the house vacuum system via Tygon tubing (∅ = 0.02 inches). The central region of this layer is comprised an array of 200□µm cylindrical micropillars, designed to evenly distribute suction force and ensure stable attachment when mated to the delivery layer. The vacuum layer featured 16 open inlets for horizontal delivery and 32 for vertical delivery, matching the number of microchannels used for each barcode set. As mentioned, the deterministic loading approach was adaptable to reagent delivery for on-chip PCR. When used for PCR, a PCR reagent-delivery layer was used and contained both an integrated network of microchannels and a microwell array matched to that of the microwell array layer. The PCR layer was designed with a single inlet for uniform and simultaneous delivery of all PCR reaction reagents and minimizing reagent loss.

### Patterning of aqueous barcodes using the deterministic barcoding technique

To assess the capacity for scale up, we sought to understand the relevant time scales for each step. A series of steps are initiated to load the barcode solutions, including: (i) placement of aliquots on inlets (∼1 min), (ii) flow into microchannels (∼ 15-20 min), and (iii) fill of microchannels with air (∼ 20 min). Next, we considered the time it takes to remove the first delivery layer and attach the second delivery layer, including alignment (∼ 10 min). We also considered loading uniformity. To assess the loading uniformity of the barcode solutions for all microwells, solutions of fluorescently labeled oligonucleotide (biotin-i51-488 and oligo-A549) were introduced into the microsystem (Figure 3D), and the coefficient of variation (CV) of fluorescence intensity in the microwells was analyzed as a measure of filling uniformity. In the horizontal loading step (Figure 3 A, B, C), the CV of the fluorescence intensity was inversely related to the loading volume (V = 0.4 µL, CV = 37%; V = 0.5 µL, CV = 29%; V = 0.6 µL, CV = 20%) until the loaded volume of 0.7 µL (CV = 24%). The slight rise in CV from 0.6 to 0.7 µL suggests microwell overfilling. The analysis indicates 0.6 µL is an optimal loading volume for this device and loading protocol design.

**Figure 3.**
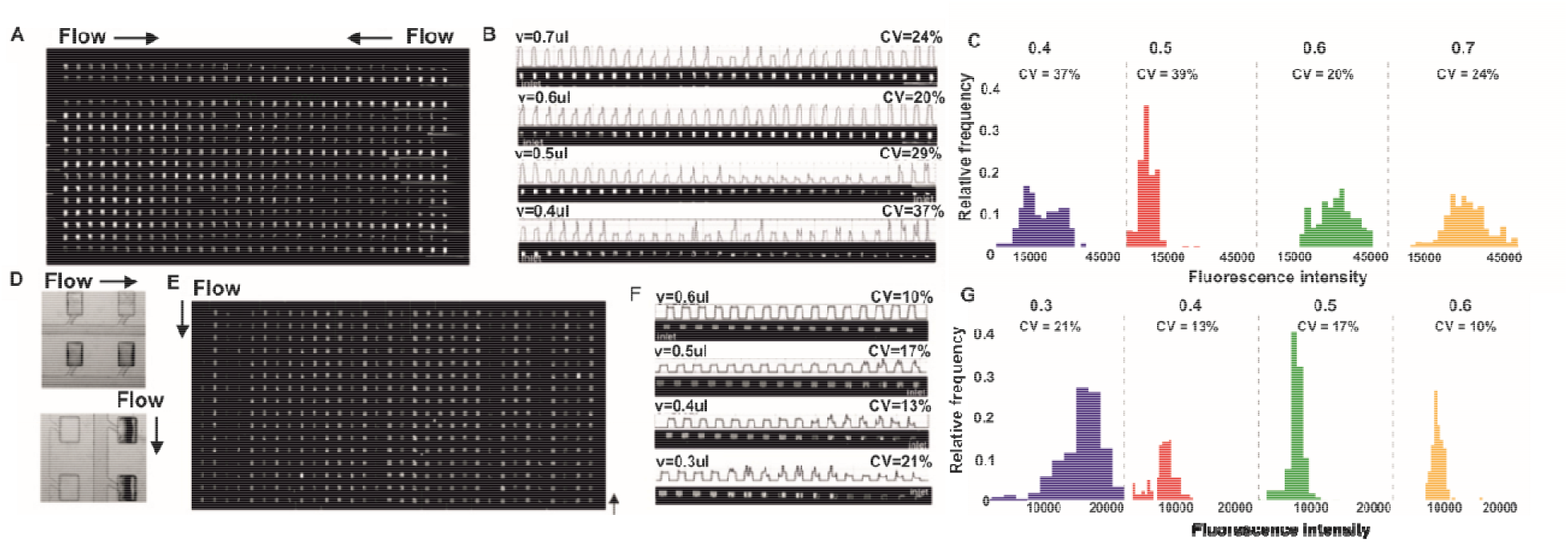
Fluorescence-based analysis of microwell loading efficiency as a function of oligonucleotide solution volume. (A) Fluorescence micrograph reporting endpoint fluorescence signal from the horizontal loading step of the aqueous barcoding workflow. Arrows who the direction of flow. (B, C) Fluorescence intensity profiles and histograms for different volumes of barcode solution loaded through the horizontal delivery layer, highlighting differences in microwell filling efficiency depending on the volume. (D) Brightfield micrographs depicting flow microchannels and microwell structures, where dark microwells indicate filled microwells and bright microwells remain empty. (E) Fluorescence micrograph reporting endpoint fluorescence signal from the vertical loading step of the aqueous barcoding workflow. Arrows who the direction of flow. (F, G) Fluorescence intensity profiles and histograms for different volumes of barcode solution loaded through the vertical delivery layer, highlighting differences in microwell filling efficiency depending on the volume.

In the vertical loading step (Figure 3 E, F, G), a similar analysis suggests the CV in microwell-to-microwell fluorescence intensity is minimized by loading a volume of 0.5 µL, with lower overall CV values for the volumes analyzed (V = 0.3 µL, CV = 21%; V = 0.4 µL, CV = 13%; V = 0.5 µL, CV = 17%; V = 0.6 µL, CV = 10%;) (Figure 3 D, E). The lower CVs for the vertical loading step versus the horizontal loading step are attributed to the lower number of microwell connected to each distribution microchannel in the vertical configuration, which likely minimizes flow resistance variability and enhances control of fluid flow. Importantly, benchmarking of the microwell loading variability of this system against similar vacuum-assisted delivery techniques^24,35,37,38^ developed for other reagent-delivery purposes is favorable.

### Cross-contamination of the deterministic barcoding technique

To evaluate the potential for cross-contamination between adjacent microwells, we scrutinized both the horizontal and vertical loading steps by imaging fluorescently labeled oligonucleotide probe signal. First, to understand cross-contamination associated with the horizontal delivery layer, we used the vacuum-assisted system to introduce a fluorescent barcode solution (oligo-A549) into every second microchannel, while concurrently filling interdigitated microchannels with a non-fluorescing solution (DI water). Fluorescence micrographs (Figure 4A) show negligible fluorescence signal in the rows of water-filled microwells, after the horizontal delivery was completed. To the same microwell array, we used a vertical delivery layer to introduce a second barcode (biotin-i51-488), which fluoresces in a different spectral channel.

**Figure 4.**
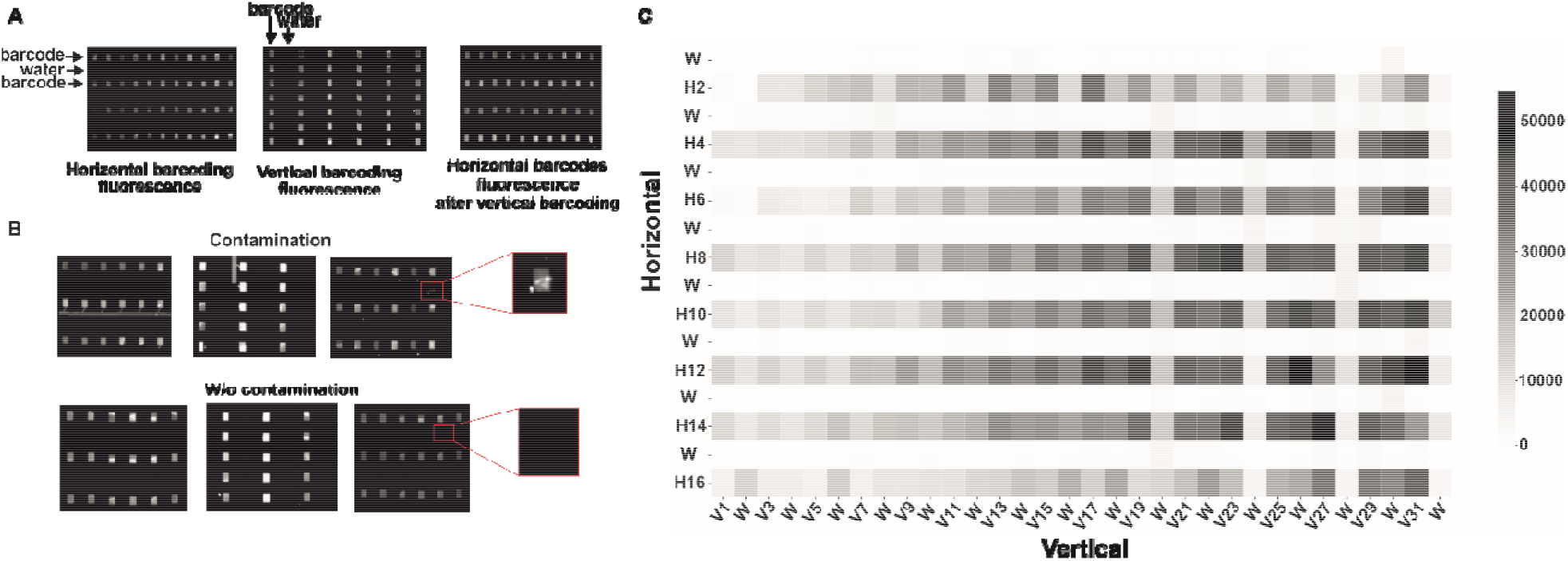
Fluorescence micrographs and a fluorescence intensity heatmap illustrating horizontal and vertical barcoding and the detection of cross-contamination between adjacent wells after the layers transition. (A) Fluorescence micrographs showing horizontal barcoding, vertical barcoding, and the resulting horizontal barcode fluorescence after vertical barcoding. (B) Representative images demonstrating the effect of contamination (top) and the absence of contamination (bottom), with insets highlighting individual microwells. Contamination (barcode transfer between adjacent microwells) can occur during layer changes (from horizontal to vertical). Contaminated microwells are marked as shown in the top panel. (C) Heatmap of fluorescence intensities corresponding to horizontal (H1–H16) and vertical (V1– V32) barcodes. In this configuration, every second row and column contained either barcode solution or water (e.g., H1 – water, H2 – barcode, H3 – water). High fluorescence intensities observed in rows or columns corresponding to water indicate contamination in those microwells.

Here, we observed one failure mode. The fluorescence micrographs in Figure 4B (top) show water-filled microwells that exhibit detectable oligo-A549 fluorescence signal, thus indicating cross-contamination introduced when swapping out the horizontal delivery layer for the vertical delivery layer. For comparison, Figure 4B (bottom) shows other microwells on the same array with no detectable cross-contamination.

Encompassing the footprint of the entire chip, Figure 4C is a heat map of fluorescence intensity from aqueous barcodes (488-nm channel) delivered by the horizontal delivery layer after completion of the two-stage barcoding workflow using the vacuum-assisted deterministic barcoding device. While detecting fluorescence signal in the horizontal barcode rows (H1-H16), we observed detectable fluorescence signal in 20 of microwells from the water-filled (W) rows, indicating ∼4.0% cross-contamination, which compares favorably to 5-10% cross-contamination for previous methods.^39–41^

### PCR-on-chip

To evaluate the performance of the deterministic barcoding strategy, we applied it to load i5 and i7 indexes into microwells for the amplification of labeled DNA, serving as the final step prior to ATAC-seq library preparation (Figure 5 A). This process was carried out following the protocol detailed in the Experimental section (PCR-on-chip). Labeling of MCF-7 nuclei with Tn5 produced an initial library concentration of 0.454 ng/µl. Subsequently, the Index-PCR master mix containing the labeled DNA was added to each microwell preloaded with dried barcodes (i5, i7). On-chip amplification for 15 cycles increased the library concentration ∼12-fold to 5.6 ng/µl, confirming efficient PCR and successful incorporation of the dual indexes across all microwells. Tapestation electropherograms (Figure 5 B) displayed the characteristic tri-modal size distribution of high-quality ATAC-seq libraries, with prominent peaks at ∼200 bp (nucleosome-free regions), ∼350 bp (mononucleosomes), and ∼550 bp (dinucleosomes). The absence of adapter dimers and the clear separation between peaks indicate high library complexity and minimal over-amplification, validating the suitability of the preparation for downstream pooling and high-depth sequencing.

**Figure 5.**
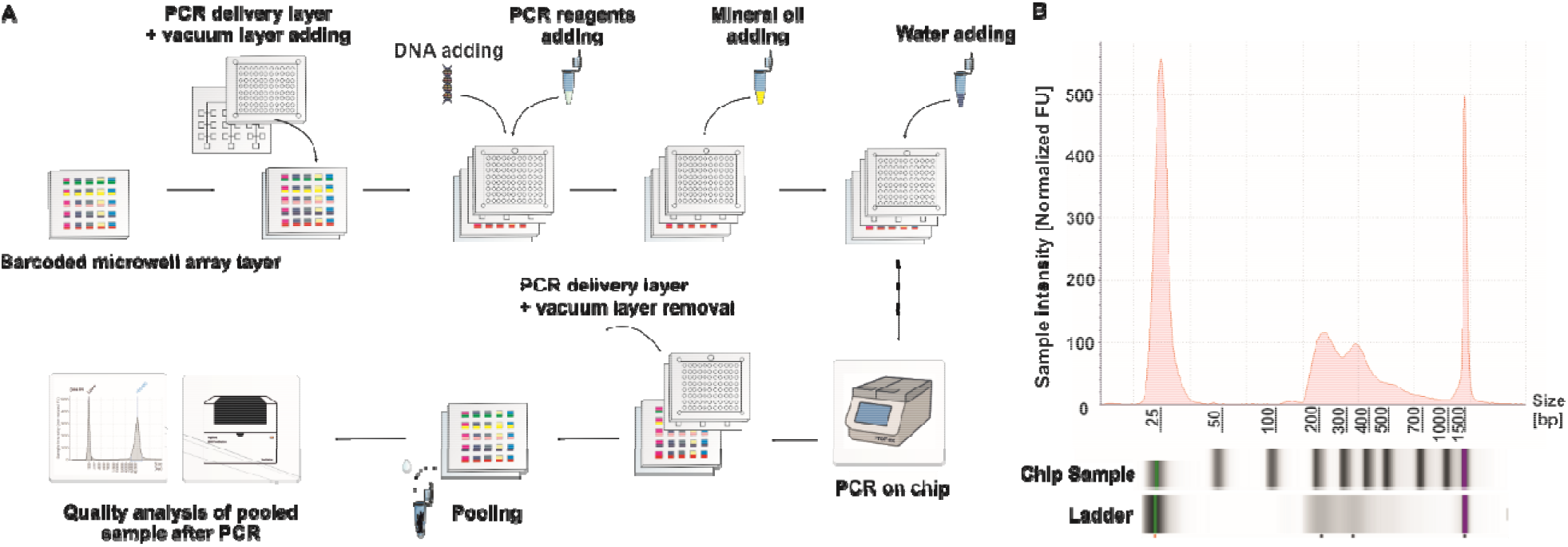
Schematic overview of the PCR-on-chip workflow and TapeStation analysis of pooled barcoded DNA samples confirming barcode integrity and fragment size distribution. (A) Schematic overview of the PCR-on-chip process. The encoded microwell array was combined with PCR and vacuum layers, enabling compartmentalized amplification after sequential addition of DNA and reagents. Pooled samples were then collected for analysis. (B) TapeStation analysis of pooled barcoded DNA samples, confirming barcode integrity and fragment size distribution.

The microwell-based deterministic indexing strategy offers several advantages over conventional tube-based workflows or previously reported microwell-based methods.^7,13,23,42^ First, pre-loading dried i5/i7 primers eliminates liquid-handling steps that can introduce cross-contamination, thereby maintaining single-sample integrity throughout amplification. Second, the confined reaction volumes (<200 nL) enhance primer□template hybridization kinetics, reducing the number of PCR cycles required to reach sequencing-ready concentrations.

## Conclusions

Barcoding strategies described in the literature are largely based on oligonucleotide-coated magnetic particles, typically delivered by peristaltic pumps.^1,43,44^ In contrast, we developed a multi-layer PDMS system in which fluid flow is driven exclusively by a standard laboratory vacuum (house vacuum), which eliminates the need for external pumps, simplifying system integration and making the platform accessible to laboratories without specialized fluidic equipment. Our platform delivers barcodes as free oligonucleotides, eliminating the need for time-consuming synthesis, preparation, and purification steps required for beads. This approach eliminates issues related to uneven bead distribution and incomplete microwell filling, while also minimizing the risk of mechanical contamination and carryover from bead surfaces.

The device features a 512-well layer, thin PDMS delivery layers, and a vacuum-connected suction layer that drives continuous liquid flow, enabling efficient delivery of 512 barcodes and precise sample labeling. We validated the system using fluorescein and fluorophore-labeled oligonucleotides to ensure uniform loading, optimize reagent use, and assess cross-contamination risk, confirming high selectivity and reliability. Deterministic assignment of i5 and i7 indexes for DNA amplification demonstrated the generation of high-quality, dual-indexed ATAC-seq libraries.

Our platform combines the strengths of existing high-throughput systems while minimizing contamination, reagent use, and library preparation time. Optimized on-chip PCR prevents leakage and material loss. Its intuitive design allows easy integration into standard laboratories, offering a flexible and scalable solution for high-throughput genomic analyses, including dual-indexed ATAC-seq.

## Supporting information

Supplementary: Experimental Section

## Acknowledgement

Funding for this work was provided by the US National Institutes of Health (R01CA20301, A.E.H.) and the Chan Zuckerberg Biohub San Francisco Investigator Award (A.E.H.). We express sincere gratitude to the Kosciuszko Foundation for the generous support provided through the Exchange to the US Fellowship during the 2024/2025 academic year, which significantly contributed to the successful completion of this research and enhanced ties between our two nations, Poland and the US.

