## Supplementary: Experimental Section for "Bead-free deterministic DNA barcoding using vacuum-driven loading of aqueous oligonucleotides to microwell arrays"

**Device Fabrication**

The geometries of all the PDMS layers were designed in AutoCAD (Autodesk), and then the masks with the mapped geometries were printed by Artnet Pro, Inc. (California, USA). To obtain stamps of individual layers, the standard photolithography technique was used.

For the horizontal, vertical, and PCR delivery membrane stamps, an SU-8 3500 (Kayaku Advanced Materials) layer was applied to a silicon wafer. It was spun using a spin coater initially for 10 s at 500 rpm with an acceleration of 100 rpm, and then for 30 s at 1000 rpm with an acceleration of 300 rpm, to obtain a layer thickness of 100 μm. The wafer was soft-baked sequentially: first for 2 minutes at 65 °C, followed by 45 minutes at 95 °C, and finally for 3 minutes at 65 °C. After baking, the wafer was exposed to UV light with a dose of 385 mJ/cm², then post-baked for 1 minute at 65°C and 5 minutes at 95°C. The wafer was developed using SU-8 developer for 5 minutes on a shaker using max rpm, then IP sprayed, dried using N2 and hard baked for 20 minutes at 200°C.

For the well layer and vacuum layer stamps production, the SU-8 2100 (Kayaku Advanced Materials) layer was applied to a silicon wafer and spun initially for 10 s at 500 rpm with an acceleration of 100 rpm, then for 30 s at 1500 rpm with an acceleration of 300 rpm to obtain a layer thickness of 200 μm. The wafer was pre-baked for 7 min at 65°C, then for 42 min at 95°C, and then cooled. The wafer was irradiated with a UV dose of 315 mJ/cm² and then subjected to a post-exposure bake for 5 min at 65°C, then 14 min at 95°C. In the next step the wafer was developed using SU-8 developer for 16 minutes on a shaker using max rpm, then IP sprayed, dried using N2 and hard baked for 20 minutes at 200°C. Next, previously prepared and degassed PDMS Sylgard 184 (Dow Corning, Midland, MI) in a 10:1 (elastomer:curing agent) was poured onto the prepared stamps. For the wells layer and all three membrane layers, the PDMS was subsequently spin-coated: first for 10 seconds at 500 rpm with an acceleration of 100 rpm, then for 30 seconds at 1000 rpm with an acceleration of 300 rpm, to achieve a PDMS layer thickness of approximately 100 μm. Next, PDMS was cross-linked by curing in an oven for 2 hours at 80°C. For the vacuum layer specifically, approximately 30 g of PDMS was poured onto the stamp, forming a layer about 1 cm thick, and subsequently cured under the same conditions (2 h at 80°C).

**Assembly of the multi-layer device and fluid routing for deterministic barcode loading**

First, the microfluidic system was prepared, consisting of a square glass microscope slide (using as a stabilizing base layer), a PDMS layer with 512 microwells, a delivery layer (horizontal, vertical or PCR depending on the test), and a vacuum layer. The glass slide was thoroughly cleaned with isopropyl alcohol (IPA, Sigma-Aldrich), rinsed with a nitrogen stream ($N_2$), and subsequently the PDMS microwell-array layer was aligned and placed on it, ensuring the removal of any air bubbles. Next, the PDMS microwell layer was treated in an air plasma cleaner (PDC-32G, Harris Plasma) at a vacuum level of 0.470 Torr using an ICME pump, with the RF power set to High for 3 min. Subsequently, depending on the type of experiment (vertical loading, horizontal loading, or PCR loading), the microwell layer, delivery layer, and vacuum layer were aligned. Plasma activation combined with vacuum suction formed a temporary, leak-free seal between the layers, allowing both reliable operation and easy layer delamination and exchange for subsequent steps. All open inlets (16 for horizontal, 32 for vertical, and 1 for PCR membrane) were sealed with tape (600 Scotch Crystal, 3M), and vacuum was applied for ~1 min. During the vacuum application, the perimeter of the microwells was visually monitored for curvature (bowing) indicating evacuation of air from the microwells and resultant vacuum-force induced distortion of the microwell perimeter at the seal interface. Once bowing was visually observed, all inlets were opened and the barcode solution (biotin-i51-488, 25 uM in ultrapure water) was introduced to the inlets using a 20 μm pipette.
